# WarpSTR: Determining tandem repeat lengths using raw nanopore signals

**DOI:** 10.1101/2022.11.05.515275

**Authors:** Jozef Sitarčík, Tomáš Vinař, Broňa Brejová, Werner Krampl, Jaroslav Budiš, Ján Radvánszky, Mária Lucká

## Abstract

**Motivation:** Short tandem repeats (STRs) are regions of a genome containing many consecutive copies of the same short motif, possibly with small variations. Analysis of STRs has many clinical uses, but is limited by technology mainly due to STRs surpassing the used read length. Nanopore sequencing, as one of long read sequencing technologies, produces very long reads, thus offering more possibilities to study and analyze STRs. Basecalling of nanopore reads is however particularly unreliable in repeating regions, and therefore direct analysis from raw nanopore data is required.

**Results:** Here we present WarpSTR, a novel method for characterizing both simple and complex tandem repeats directly from raw nanopore signals using a finite-state automaton and a search algorithm analogous to dynamic time warping. By applying this approach to determine the lengths of 241 STRs, we demonstrate that our approach decreases the mean absolute error of the STR length estimate compared to basecalling and STRique.

**Availability:** WarpSTR is freely available at https://github.com/fmfi-compbio/warpstr

## 1 Introduction

We present a novel approach for length determination of short tandem repeats (STRs) in a genome from the raw signal of nanopore sequencing reads. Our method models an STR locus using a finite-state automaton and uses an adapted version of the dynamic time warping algorithm (DTW) (Bellman and Kalaba, 1959; Senin, 2008) to find an accurate sequence of the read within the STR locus.

STRs are repetitive genomic elements containing many consecutive copies of the same short motif, typically of length 1-6 bp. The number of consecutive repeats often varies among individuals, and is one of the largest sources of intraspecies genetic diversity (Gelfand *et al*., 2006; Gymrek et al., 2016). STRs are involved in determination of quantitative traits and complex multifactorial diseases, and more than 50 STR loci in the human genome have been unambiguously associated with severe human monogenic diseases (repeat expansion disorders; REDs). REDs are typically caused by repeat length expansions of certain STRs over a pathogenicity threshold (Depienne and Mandel, 2021; Gymrek, 2017). Accurate determination of STR lengths is thus critical to differentiate between normal range, premutation range (having intergenerationally unstable lengths but without clinical symptoms), and pathogenic range alleles, but also because in several REDs, the STR length correlates with disease onset and severity (Liu *et al*., 2020; Doyu et al., 1992).

Up until recently, the STR lengths were estimated mostly by conventional methods of molecular-biology (i.e. Southern blotting and PCR-based techniques). In general, their use is complicated by the highly variable range of allele lengths to be detected, by the immense length of the expanded ones, and by the stable secondary structures formed by these repeats. Moreover, the accuracy of these methods decreases significantly with the length and complexity of the STR motif. Therefore, although several improvements have been introduced to overcome these challenges, there is still no single method that would reliably identify and size all possible ranges of normal and expanded alleles, and it is recommended to use them in different parallel combinations. These methods are however limited to characterisation of individual loci and are not suitable for genome-scale screening (Bahlo *et al*., 2018; Radvansky and Kadasi, 2010). In contrast, short-read sequencing allows to study STRs in more detail and on the whole-genome scale. However, properly genotyping STRs of lengths surpassing the used read length is difficult and at best limited to the identification of the presence of such alleles, but not their size (Budiš *et al*., 2018; Dashnow *et al*., 2018; Willems *et al*., 2017; Halman and Oshlack, 2020; Dolzhenko et al., 2019).

Recently, long-read technologies (such as Pacific Biosciences, and Oxford Nanopore) offered new possibilities for studying even larger expanded STRs (Sedlazeck *et al*., 2018). However, higher sequencing error rates pose additional challenges. One of the first tools using long reads, PacmonSTR (Ummat and Bashir, 2014), modifies a traditional alignment algorithm to take higher error rates of Pacific Biosciences reads into account. The resulting estimate of the repeat number is subsequently corrected by a pair hidden Markov model. RepeatHMM (Liu *et al*., 2017) performs error correction of repeat sequences in Oxford Nanopore reads by aligning them with the template of perfect repeats of the specified pattern using asymmetric sequence alignment. Then, RepeatHMM uses a hidden Markov model (HMM) to estimate the number of repeats directly from the corrected read sequences. In both tools, the results from individual reads are pooled together using Gaussian mixture models (GMMs) to predict the final genotype. TandemGenotypes (Mitsuhashi *et al*., 2019) uses a tailored alignment strategy based on last-split (Frith and Kawaguchi, 2015) to obtain more confident alignments between the reference repeat region and reads. The change of the repeat number present in the read compared to the reference is then computed as a sum of contributions from the number of unaligned bases in the reference and the read.

Other tools, such as NanoSatellite (De Roeck *et al*., 2019) and STRique (Giesselmann *et al*., 2019), attempt to overcome high error rates in nanopore sequences by using the raw sequencing signal. The raw signal is comprised of measurements of electrical current as DNA passes through a nanopore. The signal is affected by a context of *k* nucleotides (typically *k* = 6) and each context is read approx. 8-9 times on average. The portion of the signal corresponding to the shift by one base is called an *event*. Typically, the raw signal is first translated into the DNA sequence by a basecalling software using complex machine learning models. The above mentioned tools use the raw signal to overcome errors introduced through imperfect basecalling.

NanoSatellite (De Roeck *et al*., 2019) uses an expected signal corresponding to five copies (and later three copies) of the repeating motif to match sections of the real signal using a dynamic time-warping (DTW) algorithm (Bellman and Kalaba, 1959; Senin, 2008). The results are clustered to two clusters to obtain the final copy number call. STRique (Giesselmann *et al*., 2019) uses both flanking sequences and the repeating pattern to build a profile HMM, where match states correspond to *k*-mers in these sequences; the STRique model does not allow for variation within the repeat. The copy number is then determined by counting match states as the raw signal is aligned to the profile HMM.

In this paper, we present a new WarpSTR tool, which improves on NanoSatellite and STRique in several ways. First, we recognize that basecalling is typically much more accurate in non-repeating regions (such as sequences flanking an STR) than in regions containing short repeats, and thus it is more efficient (and even more accurate) to simply use basecalled sequences to locate the flanks and subsequently isolate the signal corresponding to the STR locus. Second, instead of using greedy heuristics as in NanoSatellite, we model the whole STR locus by a finite-state automaton and modify the DTW algorithm to align the full length of the signal corresponding to the STR locus to this automaton, determining the corresponding sequence length by the number of states passed by the alignment path. This is similar to the STRique profile HMM; however, there are several major differences. First, in WarpSTR, the finite-state automaton enables much greater customization. This allows us to model more complex STRs that include combinations of heterogeneous motifs or are interrupted, which are common occurrences in medically relevant STRs (Musova *et al*., 2009; Radvanszky *et al*., 2021; Andrew *et al*., 1994; Radvansky *et al*., 2011). Second, WarpSTR attenuates the signal normalization problem using a novel signal polishing phase. Finally, we use Bayesian Gaussian mixture models to summarize the information from multiple overlapping reads and to derive the final genotypes. Using nanopore reads from whole-genome sequencing of human genomes and comparing our results to high-confidence variant calls determined by integrative approaches, we determine that our approach is significantly more accurate than both NanoSatellite and STRique.

## 2 Methods

The speed of DNA passing through a pore varies significantly, which makes signal-based analysis challenging (Midha *et al*., 2019). To address this problem, dynamic time warping (DTW) technique (Bellman and Kalaba, 1959; Senin, 2008), which has been used e.g. in speech processing, has been adapted to nanopore signal analysis (Loose *et al*., 2016; Han et al., 2018, 2019). To align a known DNA sequence *S* to a raw signal by DTW, we first estimate an “ideal” signal observed by sequencing *S*. Namely, we decompose string *S* to a sequence of overlapping *k*-mers and for each *k*-mer use the value from the lookup table of the expected signal values provided by Oxford Nanopore Technologies (Oxford Nanopore Technologies, 2017a). This signal can be then aligned to the real signal using dynamic programming, minimizing an error function based on the difference between the two compared signal points. In contrast to sequence alignment, alignment of multiple points in one signal to one point in the other signal is not penalized by DTW due to the expected wide variation in signal speeds.

In our work, we adapt DTW for alignment of a repetitive sequence pattern of an undetermined length. Using the alignment of the known sequence pattern to the raw signal helps to inform the decisions in case of ambiguities, and ultimately results in a more accurate interpretation of the raw signal. Once the pattern, representing an STR, is aligned to the raw signal, its instantiation is fixed, and the length of the instantiation determines the length of the STR represented by the nanopore read.

### 2.1 STR Signal Extraction

WarpSTR assumes that the input data was preprocessed by basecalling the raw reads and mapping them to a reference genome using standard methods. For each read overlapping the target region, the flanking sequences are then found in the basecalled sequence using local alignment, and the mapping between the called bases and the raw signal, which is an interim output of the Guppy basecaller (Wick *et al*., 2019), is used to locate the corresponding signal positions. In the experiments, we have used Guppy v.4.0.15 and the flanking sequence length of 110 bases.

### 2.2 STR Representation by a Finite-State Automaton

An STR locus of an individual haplotype may differ from the reference genome in the number of repeats and in substitutions in individual repeat instances. To represent variability of each STR, we use a finite-state automaton as shown in Figure 1, constructed from a simple regular expression. In our software, we currently support IUPAC nucleotides, regions that can be skipped denoted by *{}*, and repeats occurring at least once denoted by (). For example, the DM2 locus, which consists of a CAGG repeat, potentially interrupted with CAGA or CAGC, can be modelled as (CAGG*{*CAGM*}*), where M is the IUPAC code for nucleotides A or C. Such simple regular expressions are sufficient to represent loci that were of interest in our analyses, however it is straightforward to extend the approach to more complex regular expressions. In the finite-state automaton, we also include flanking sequences from the reference genome.

**Figure 1:**
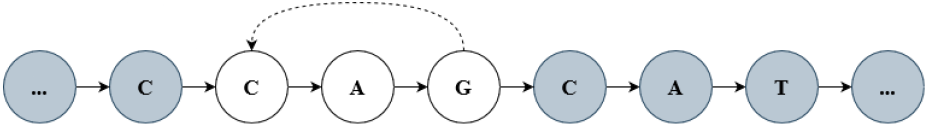
A finite-state automaton modelling a DM1 STR locus (CAG) and its flanking sequences (shown in gray).

In the next step, we extend the state space of the automaton to the *k*-mer space to keep track of the context of the last *k* nucleotides, and transform the original automaton to a new finite-state automaton over *k*-mers (see example in Figure 2). After this transformation, we can assign to each state the expected value of the nanopore signal aligned to this state.

**Figure 2:**
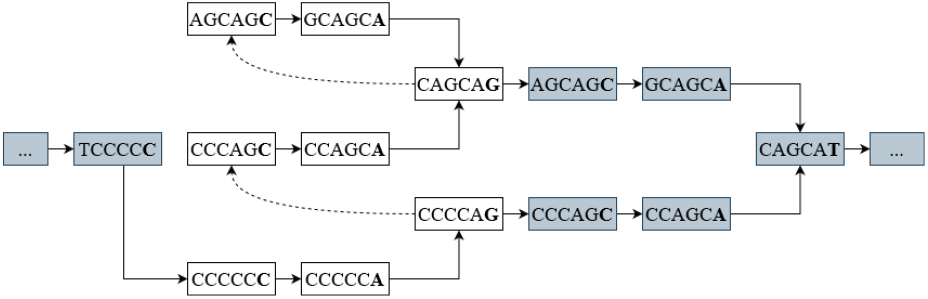
Extended finite-state automaton over the *k*-mer space

To align the raw read signal *S* = *s*_1_ … *s*_*m*_ to the finite-state automaton, we use dynamic programming analogous to DTW, where subproblem *M*_*i,j*_ represents the cost of the best alignment of the first *i* signal points *s*_1_ … *s*_*i*_ under the condition that we finish in state *j*. Values *M*_*i,j*_ can be computed using the following recurrence:

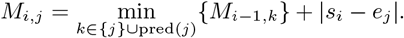

Here, pred(*j*) is the set of all predecessor *k*-mers of state *j* in the automaton and *e*_*j*_ is the expected signal level for the *k*-mer represented by state *j*. A straightforward implementation of this recurrence works in *O*(*mnd*) time, where *m* is the length of the signal, *n* is the number of states in the automaton, and *d* is the maximum size of pred(*j*). By counting the number of state transitions and subtracting the flanking sequences, we obtain the estimated length of the STR (see Figure 3 for illustration).

**Figure 3:**
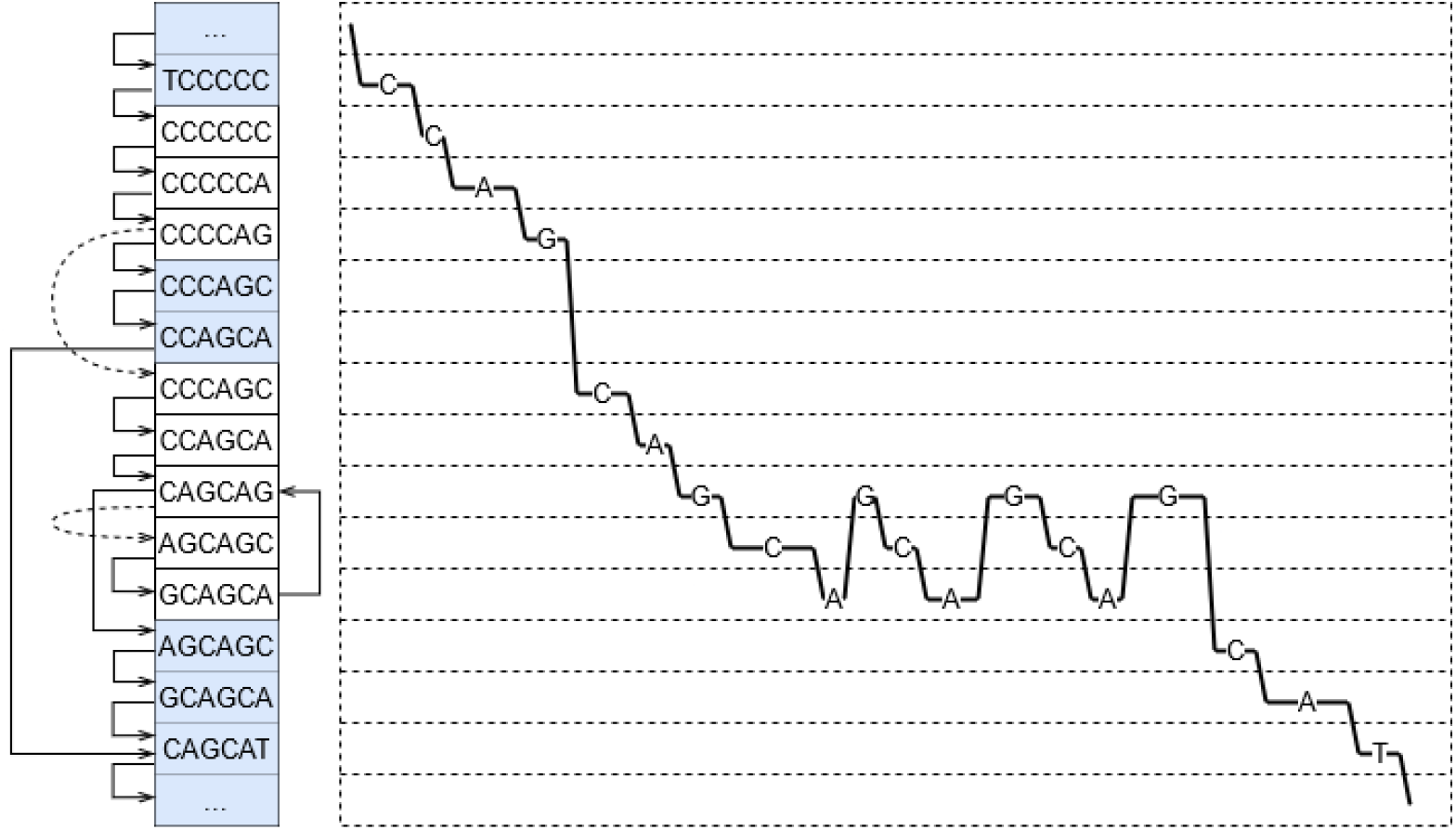
A part of the warping path produced by the WarpSTR search algorithm using the state automaton for DM1 locus. States representing *k*-mers are shown on the left, while the part of the dynamic programming matrix is shown on the right. The warping path, i.e. the path through states with the lowest cost is shown as the sequence of lines with corresponding nucleotides.

When there is no user-defined regular expression modelling the repeat, WarpSTR creates it automatically from the reference genome using an input file listing boundary positions of STR regions of interest and their repeat patterns. The software creates the expression using the exact copies of the pattern found in the reference and any incomplete copies or interruptions. For example, if the reference sequence of the STR region was GGG followed by 6 repeats of the pattern AGAGGG, the automatically generated regular expression was GGG(AGAGGG). These automatically created regular expressions can be adjusted by the user to add flexibility, for example interruptions not present in the reference genome.

### 2.3 Signal normalization

Nanopore signal is scaled and shifted differently in each sequencing read and it needs to be normalized before analysis so that the resulting values can be compared to the expected signal levels defined in the *k*-mer tables. The most common approach is based on the assumption that each read represents a sufficiently long random sequence, and thus basic statistics, such as the mean, the median, and the variance should match across different reads. We apply the median normalization strategy outlined in tombo (Oxford Nanopore Technologies, 2017b), which for a raw signal sequence *R* = *r*_1_ … *r*_*n*_ computes values *shift* = 1*/*2(perc_46.5_(*R*) + perc_53.5_(*R*)) and *scale* = median_*i*_(|*r*_*i*_ *− shift* |), where perc_*j*_(*R*) is the *j*-th percentile of values *r*_1_ … *r*_*n*_. The normalized signal *S* = *s*_1_ … *s*_*n*_ is obtained from *R* as *s*_*i*_ = (*r*_*i*_ *−* shift)*/*scale.

Characteristics of a signal produced from a long repetitive region may differ substantially from a random sequence, and consequently the values obtained by the median normalization may not match those in the expected signal level tables. This is likely one of the reasons why basecalling in these regions often exhibits large errors. To address the problem, we first use the standard normalization and our DTW algorithm to obtain an initial alignment of the raw signal to the finite-state automaton. We compute the mean *m*_*j*_ of all values aligned to a particular state *j*, and we pair each *m*_*j*_ with the corresponding expected signal *e*_*j*_ from the model. Some portions of the signal could be aligned incorrectly, and we discard all pairs (*m*_*j*_, *e*_*j*_) where *m*_*j*_ is further from the expected signal *e*_*j*_ than a given threshold.

To *polish* the signal, the set of remaining pairs is approximated by a spline using splrep and splev functions from SciPy (Jones *et al*., 01), and signal value *x* is then replaced with the spline value for this *x* coordinate, which corresponds to an interpolated expected signal (see Supplementary Section S1). The second iteration of our algorithm uses the polished signal and the final length of the STR is determined based on its output. Figure 4 illustrates that even small changes resulting from polishing can lead to changes in the subsequent second alignment and the resulting repeat count.

**Figure 4:**
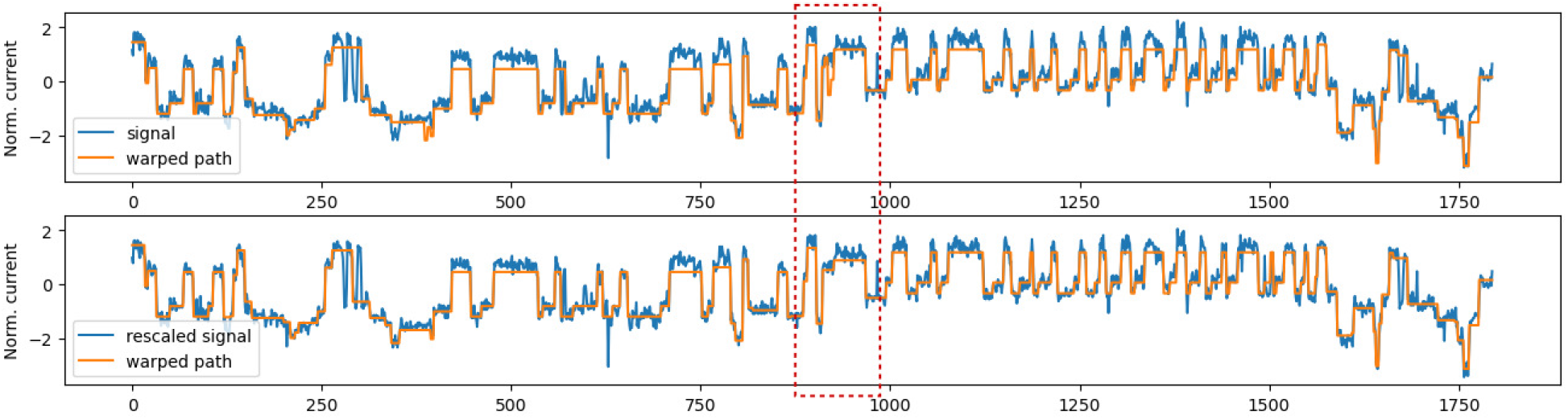
Signal polishing effect on the alignment. Example alignment of the signal to the expected signal from the state automaton before (top) and after polishing (bottom). Before polishing, the normalized signal values are much higher than the expected signal values, and some of these differences decrease after polishing. More importantly, a spurious repeat, highlighted in the red window, disappears after polishing.

### 2.4 Adaptive restriction of event size

In some cases, the expected signal levels in different states are very similar to each other, and the noise present in the raw signal may drive the algorithm to match the same real event to multiple distant states by using several very small events to get to a new state. This would overestimate the STR length. We have solved this problem by requiring a given minimum number *s* of signal points to be aligned to each state, eliminating short events altogether.

Setting this parameter is important for accuracy. Using a too large fixed value may cause short consecutive to be skipped altogether. To remediate this problem, we set the parameter adaptively, i.e. we decrease the minimum number of signal points adaptively when short consecutive events occur in the signal.

To find parts of the signal with short consecutive events, we use the result of alignment with *s* = 4. Alignment result is split into non-overlapping windows containing signal points aligned with *m* events (in experiments we used *m* = 6). Window *W* of *w* signal values is looped through in an overlapping sliding window manner with sliding window length 2*p* to obtain *w −*2*p* subwindows. Similarly as in (Zhang *et al*., 2021), for each subwindow *S* of length 2*p* signal values, Welch’s unequal variances t-test is calculated as follows: 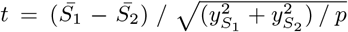, where *S*_1_ and *S*_2_ are *S*[0, *p*) and *S*[*p*, 2*p*) respectively, *y* represents the standard deviation and 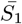 represents the mean of *S*_1_. When an extreme value *t* is calculated, it denotes that there is a significant context change in that subwindow. By simple local optima picking of *t* values from all subwindows we obtain the set of peaks, denoting the expected number of context changes in the whole window. When the window has more context changes than *m*, WarpSTR presumes that some short events were missed. Thus, in the second signal polishing phase the values of *s* is decreased by one for those particular windows.

### 2.5 Summarizing the results

After running the search algorithm, we obtain a length estimate for each read overlapping a particular STR locus. For diploid organisms, there could be two different alleles with different lengths (one inherited from each parent). Thus we want to summarize all of the estimates by either one value (if the locus is homozygous) or two values (if the locus is heterozygous).

WarpSTR first filters out apparent outliers by removing estimates that are further than two standard deviations from the mean estimate. To separate reads into two groups, we used Bayesian Gaussian mixture models with maximum of two components from the Python scikit-learn library (Pedregosa *et al*., 2011) (with settings tied covariance type, the weight concentration prior 0.25, 5 initiations, 1000 iterations). WarpSTR declares an STR locus as homozygous if the number of reads in the smaller cluster does not exceed 20% of all unfiltered reads. In this case, the median allele length is reported as the final estimate. Otherwise, the STR locus is declared as heterozygous, and the medians of the two discovered clusters are reported.

## 3 Experiments

We have used publicly available nanopore data sets for human samples NA12878 (Jain *et al*., 2018) and NA24385 (Oxford Nanopore Technologies, 2020b,a) produced with R9.4.1 nanopore flow cells. We have downloaded the raw fast5 files and BAM files produced using minimap2 and human reference genome version hg38. The NA12878 data set was used for finding the optimal settings of hyperparameters while the NA24385 data set was used for final evaluation only.

To compare to other tools, which cannot handle complex STRs, we have randomly selected 433 STR regions with a single repeating pattern of length between 2 and 6 (i.e. no homopolymeric repeats) and the reference sequence of STR region could not contain a homopolymeric sequence longer than 6. To this end, we have used information provided in HipSTR (Willems *et al*., 2017). The input regular expressions required by WarpSTR were automatically generated using the repeat pattern and the reference genome. In the second experiment, we use complex STRs to further demonstrate WarpStr capabilities.

High-confidence benchmark VCF files assembled by a complex integrative analysis from multiple sequencing data sets (Zook *et al*., 2019; Wagner et al., 2020) (v.3.3.2 for NA12878, v.4.2.1 for NA24358) were used as a gold standard. Entries in the VCF files represent predicted variations compared to the reference genome, while also showing types of variations (i.e. whether it is a substitution, insertion, or deletion) and genotype information, i.e. whether the variation is homozygous, or heterozygous, in which case could contain either one or two alternative genotypes. As used VCF files stored combined predictions of 8 different genotype callers, at first, the allele lengths of all loci were obtained for each caller. The allele lengths were computed by adding the length of insertions or subtracting the length of deletions from the reference allele length and using the information about genotype. For example in case of reference allele length of 40, and a heterozygous insertion with just one alternative allele of length 4, and a homozygous deletion of 2 nucleotides, the final allele lengths were given as 38,42. To discard spurious or low confidence allele length estimations, caller estimations were combined as follows. If only one caller had allele length estimation, then it was discarded. When two callers had allele length estimations and both of them were equal, then the estimation was taken as the true answer. When *k* ≥ 3 callers had estimations and *k −* 1 of these callers had the same estimation, then it was taken as the true answer. In other cases or when there were no entries in the VCF file for a locus, the locus was discarded. This resulted in a final set of 241 loci with ground truth information.

### 3.1 Obtaining predictions for comparison

As a baseline prediction, we have obtained allele lengths directly from basecalled reads, by aligning the left and right flanking sequences of length 110 with each read. If the alignment score was not high enough or if the right flank was found before the left flank, the read was filtered out. The predicted allele lengths were further processed by filtering and clustering in the same manner as in WarpSTR, to obtain the final result.

We compare our results to STRique (Giesselmann *et al*., 2019)(v0.4.2.) with default parameters. The length of flanks was set to 150 as recommended by STRique authors. We discarded reads where the repeat region was incorrectly extracted by STRique or with the weak signal alignment scores for flanks (less than 3.8) or where the resulting prediction was 0.

### 3.2 Results

First, we compared STR length estimates for individual reads using the mean absolute error (MAE) and median absolute error (MedAE). For length calls *y*_1_, …, *y*_*n*_ and the gold standard answer (*m*_1_, *m*_2_), these measures are defined as MAE = avg_i_*{*min*{*|*y*_*i*_ *− m*_1_|, |*y*_*i*_ *− m*_2_|*}* and MedAE = median_i_*{*min*{*|*y*_*i*_ *− m*_1_|, |*y*_*i*_ *− m*_2_|*}*. The results are shown in Table 1.

**Table 1:** The performance comparison of WarpSTR, STRique and the baseline on 241 loci.

|  | WarpSTR | baseline | STRique |
| --- | --- | --- | --- |
| Discarded reads | 10.75% | 10.75% | 26.81% |
| # loci with the lowest MAE | 204 | 36 | 0 |
| # loci with the lowest MedAE | 237 | 152 | 64 |
| Global MAE | 3.84 | 4.89 | 334.8 |
| Global MedAE | 0 | 1 | 4 |
| Average D | 0.73 | 0.96 | 11.74 |
| Median D | 0 | 0 | 6 |
| # of loci with D=0 | 205 | 182 | 37 |
| Running time - 1 thread | 345.43 | - | 926.08 |
| Running time - 2 threads | 196.79 | - | 481.83 |
| Running time - 4 threads | 123.52 | - | 274.1 |
| Running time - 8 threads | 106.47 | - | 256.72 |

WarpSTR produced estimates with the lowest MAE in 204 out of 241 loci. STRique performs poorly under this measure due to outliers occurring frequently even after score filtering recommended by the authors (we assume that this may be due to imprecise signal extraction in STRique). MedAE (median-based measure) is less prone to such outliers, but it created many ties between the tools. However, WarpSTR still performed the best in almost all cases. Supplementary Figure S2 illustrates the distribution of MedAE across loci. When we combine predictions for reads from all loci to a single global MAE and MedAE score, WarpSTR also performs the best. WarpSTR uses data more efficiently than STRique (only 11% of reads were discarded due to inaccurate alignment of flanking sequences in WarpSTR, while STRique discards more than 26%).

Interestingly, the MAE for individual loci is influenced by the repeated pattern length (see Figure 5). The patterns of length two are in general the most difficult for both methods. In such case, the expected signal has only two alternating values, a pattern easily confused with noise. Even in these cases, WarpSTR outperforms basecalling in approximately half of the loci. For longer patterns, the error is generally lower and WarpSTR very consistently produces the best results.

**Figure 5:**
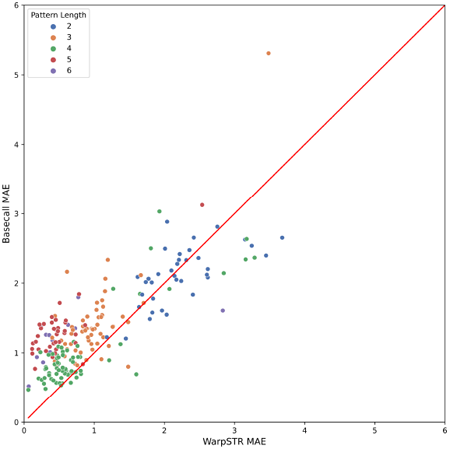
MAE for WarpSTR and basecalling for individual loci colored by repeating pattern length. 25 loci that have very high MAE in both methods are not shown; these are most likely due to large expansions not captured by VCF callers in the gold standard.

So far we have compared predictions for individual reads. These were then combined to an overall genotype for each tool using the filtering and clustering procedure described in the Methods section (the same method was used for all tools). Let *ŷ*_1_, *ŷ*_2_ be the final predicted allele lengths and *y*_1_, *y*_2_ the true alleles, assuming *ŷ*_1_ ≤ *ŷ*_2_, *m*_1_ ≤ *m*_2_. As the error metric, we use *D* = |*ŷ*_1_ *− m*_1_| + |*ŷ*_2_ *− m*_2_| if the prediction or the true answer or both are heterozygous, and *D* = |*ŷ*_1_ *− m*_1_| if both are homozygous. As we can see in Table 1, WarpSTR performs better than other tools, providing perfect results in 85% of the loci (205 out of 241).

Table 1 also shows a comparison of the running time. As the tools have very different preprocessing and postprocessing steps, we only compare the core part of the pipelines (see Supplementary Section S3). WarpSTR is generally faster than STRique; apart from implementation differences, the main reason is that WarpSTR uses only matching states in the model, whereas STRique also uses insertion and deletion states.

One of the advantages of WarpSTR is its ability to analyze STRs with a complex structure, compared to other methods, which are limited by a single repeat pattern. We will illustrate this strength on two clinically relevant complex STR loci: HD (Huntington’s disease) (Andrew *et al*., 1994) and DM2 (Myotonic dystrophy type 2) (Radvanszky *et al*., 2021).

The HD locus consists of AGC and CGC repeats separated by a four codon sequence AACAGCCGCCAC, which is not prone to repeat. Thus, the input sequence for WarpSTR was (AGC)AACAGCCGCCAC(CGC). For the reads included in the NA24385 sample, the WarpSTR predicted alleles of lengths 84 and 105, which agreed with the gold-standard answer. Basecalling prediction was 83 and 102 (see Supplementary Section S4). STRique cannot be run on this locus, as it works only for simple repeat patterns.

Thanks to the usage of a state automaton, it is possible to count the number of occurrences of each repeating part of the input sequence, as given by parentheses in the WarpSTR input sequence, and these can be further clustered into genotypes. In the gold standard, HD has 17 and 24 repeats of AGC, and 12 repeats of CGC, and using WarpSTR, we came to the same result (see Supplementary Section S5).

The DM2 locus is particularly complex, consisting of CAGG repeats with CAGA or CAGC interruptions, followed by CAGA repeats and finally CA repeats. The input sequence for WarpSTR was given as ((CAGG)CAGM)(CAGA)(CA), where CAGM denotes both CAGA and CAGC interruptions. Figure 6 shows clustered predictions of DM2 for NA24385 subject split per repeat unit, also listing counts of individual interruptions. The first allele was predicted to contain 16 CAGG repeats of which 2 are CAGA interruptions and 1 CAGC interruption, followed by 8 CAGA repeats and 21 CA repeats. The second allele was predicted as 18 CAGG repeats of which 2 are CAGA interruptions and 1 CAGC interruption, followed by 6 CAGA repeats and 25 CA repeats.

**Figure 6:**
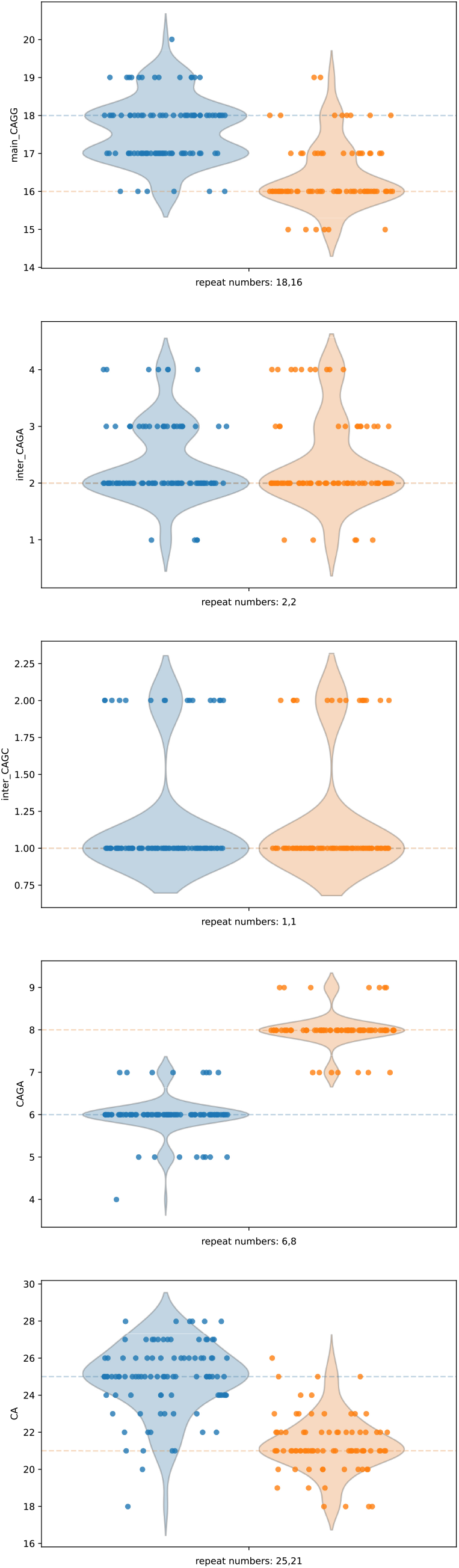
Clustered predictions of DM2 f9or NA24385 subject split per repeat unit.

## 4 Conclusion

In this paper, we have investigated the problem of estimating the lengths of short tandem repeats (STRs) from nanopore sequencing reads. Nanopore sequencing is a promising technology, providing very long sequencing reads at a reasonable cost and throughput (a single Promethion flowcell can sequence a human genome at a high depth). Long read sequences provide a unique opportunity to study the role of STRs in various diseases, since previously employed short-read technology has problems with genotyping larger loci that can be even several hundred bases long.

Since basecalled reads from nanopore sequencing typically exhibit lower accuracy in STR regions, sometimes to the point at which the repeats are unrecognizable, we opted for analyzing the raw sequencing signal instead. We have adapted a commonly used dynamic time warping technique to work with representations of the STRs based on a finite-state automaton and determined the length of an STR by alignment of the raw signal to such an automaton. Proper scaling and shifting of the raw signal is also difficult in regions with repetitive sequences. To address this problem, we have developed a new method for polishing the raw signal using splines. Another innovation is an adaptive setting of the minimum event size for parts of the raw signal. The resulting software tool called WarpSTR is able to genotype STR alleles with high accuracy while outperforming baseline approach employing the basecalled sequences and STRique, another tool recently developed for this purpose.

One obvious extension would be to enrich the underlying finite-state automaton to allow for previously unmapped insertions, deletions, and substitutions. This would likely require employment of a probability-based scoring scheme instead of the simple scoring scheme used in the present work. In fact, defining a probabilistic model for the problem would allow us to use more advanced techniques to deal with high levels of uncertainty in nanopore raw signals, and perhaps allow us to predict a posterior distribution of STR length instead of a single value that can potentially harbor systematic errors.

Finally, at present we use a simple clustering scheme to summarize the results as a genotyping call, but perhaps techniques modeling typical errors observed in STR analysis would lead to a further improvement.

