## Supplementary materials for "WarpSTR: Determining tandem repeat lengths using raw nanopore signals"

#### S1 Signal polishing

Supplementary Figure S1 shows the polishing effect of state signal values compared to the expected signal values. A spline is computed from the sorted mean signal values for each state (in blue) and the expected signal value for the corresponding state (in green). Using spline, signal values are polished and new state means (in orange) are obtained (just for visualization purposes). We can see in the figure, that some new state means are closer to the expected means, possibly reducing the distance between the signal values and expected signal values, therefore obtaining more accurate alignment in the second phase.

#### S2 High similarity of event values

It is common knowledge that homopolymeric sequences are impossible to characterize correctly using nanopore sequencing as the measured signal is just a long stretch of near-constant signal values with some length, but due to the variable strand velocity, it is not possible to infer the homopolymeric sequence length.

Very low variance signal values can also occur when some repeating pattern produces very similar signal values for each possible k-mer. This is problematic for basecalling as for WarpSTR or any other tool. A good example is the pattern AAGAG, which repeats on the reverse strand in a genomic region given by coordinates chr10:8332996-8333065 on the hg38 reference genome (see Figure S3).

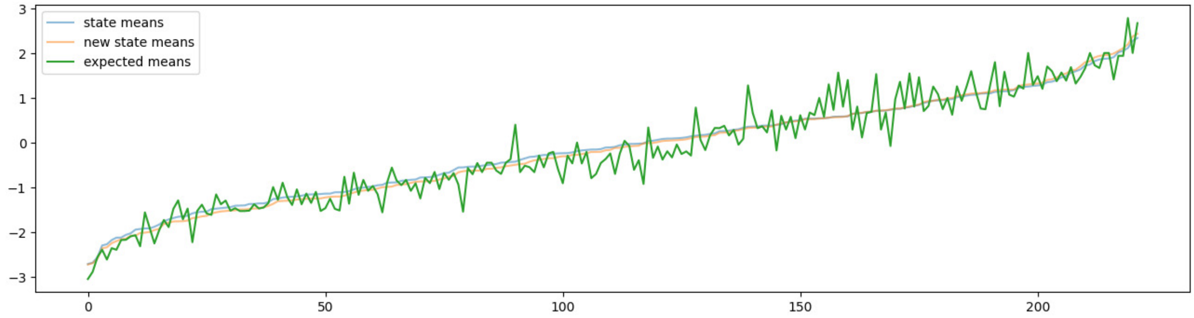


**Supplementary Figure S1:** Signal polishing using spline.

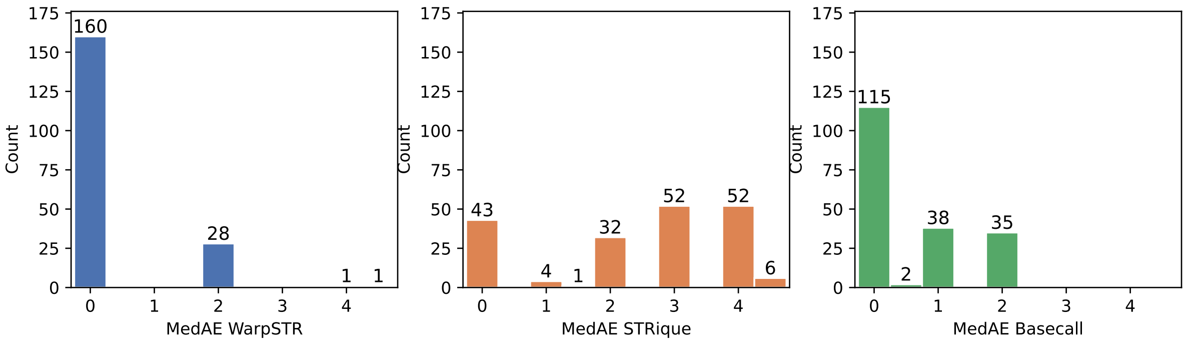


**Supplementary Figure S2:** The histograms of MedAE per locus for individual tools, capped at 5.

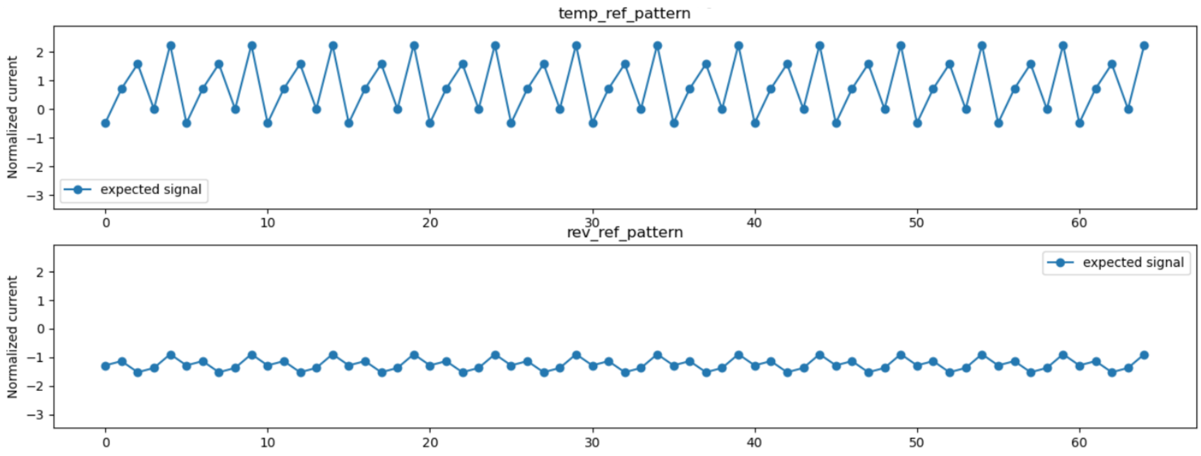


**Supplementary Figure S3:** The expected signal of repeats of pattern CTCTT and AAGAG (on the reverse strand).

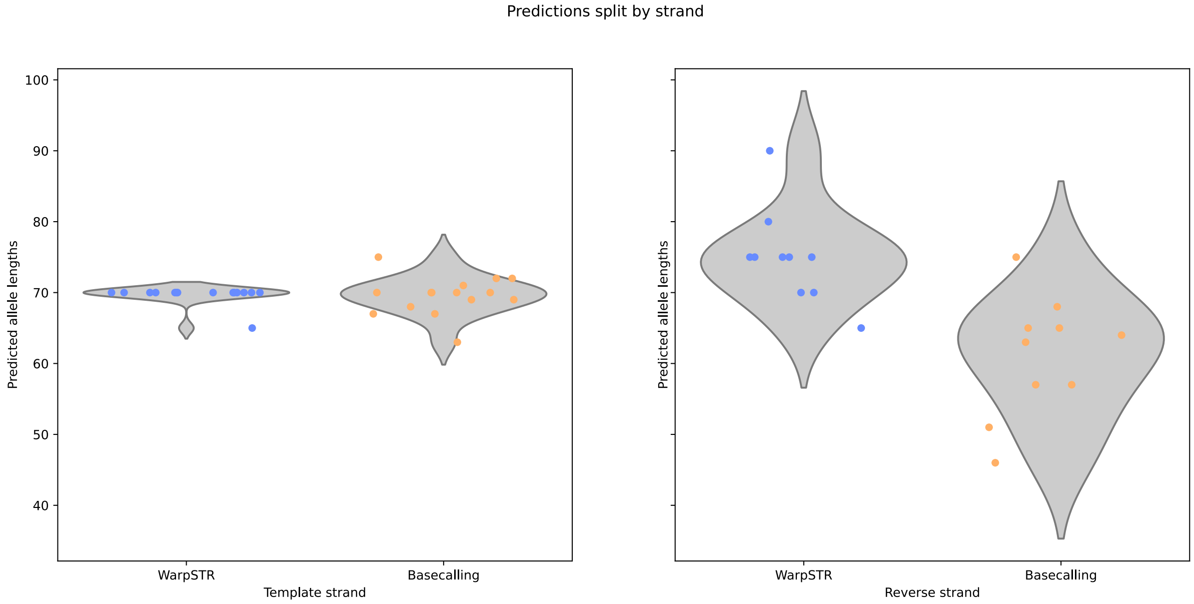


**Supplementary Figure S4:** Predictions for a repeat of pattern with low (on template strand) and high inter-state similarity (on reverse strand).

Supplementary Figure S3 clearly shows that the events of the repeating signal on the reverse strand are very similar to each other in their normalized expected signal values, while on the template strand, this does not hold, and the events are very well distinguishable apart from each other. This usually creates a situation where predictions on the high-similarity strand have much larger error (see Figure S4).

In WarpSTR, we warn users about this problem by computing the mean absolute difference of consecutive signal values. In this example, mean state difference was 1.727 and 0.312 for template and reverse strand, respectively.

Supplementary Figure S4 shows predictions split by strand, using NA12878 data, where the number of repeats as given by the golden set is 14, i.e. allele length is 70, for both alleles. The figure also shows how this high state similarity influences basecalling accuracy. On the left, the figure shows violin plots for allele lengths as given by WarpSTR and basecalling for the template strand, and for the reverse strand on the right. We can see that the problem of low variance signal is difficult to overcome also for basecallers.

Using this mean state difference as given by WarpSTR, we can prevent incorrect determination of repeat length due to the low variance in repeating pattern on one strand. However, it is also important to note, that in cases where both strands have very high similarity, this does not give a powerful message apart from the information, that the prediction will be of lower confidence for that locus.

### S3 Details of running time measurements

As both tools have different configurations and initial steps, we have measured only the time of finding the STR signal part and determining the repeat count. For STRique, it was the running time used for calling the ‘count‘ option of STRique tool. For WarpSTR, the running time was defined as the time required to extract the repeating signal, generate expected signals, construct the automaton, and run both alignment phases. As clustering is not implemented in STRique, WarpSTR clustering is not measured. Parameters were the same for WarpSTR and STRique as in other experiments.

For this experiment, we used a personal laptop with an AMD Ryzen 7 4800H processor and Ubuntu 20.04 as OS. First, we chose 10 random loci and obtained reads for them for NA12878 subject with 20-30 reads for locus. Then we used STRique and WarpSTR 5 times and measured the running time, which we averaged in the end. The experiment was run using 1, 2, 4 and 8 threads.

### S4 Per locus predictions

WarpSTR also visualizes (if set in configuration file) the distribution of WarpSTR and basecalling predictions, with the final predicted alelle length noted under the plot.

Supplementary Figure S5 shows such results for HD locus of NA24385 subject. Gold standard shown two alleles, one with 84bp and the other with 105bp. WarpSTR achieved the same result, while basecalling shown a small error while it reported alelles with 83bp and 102bp.

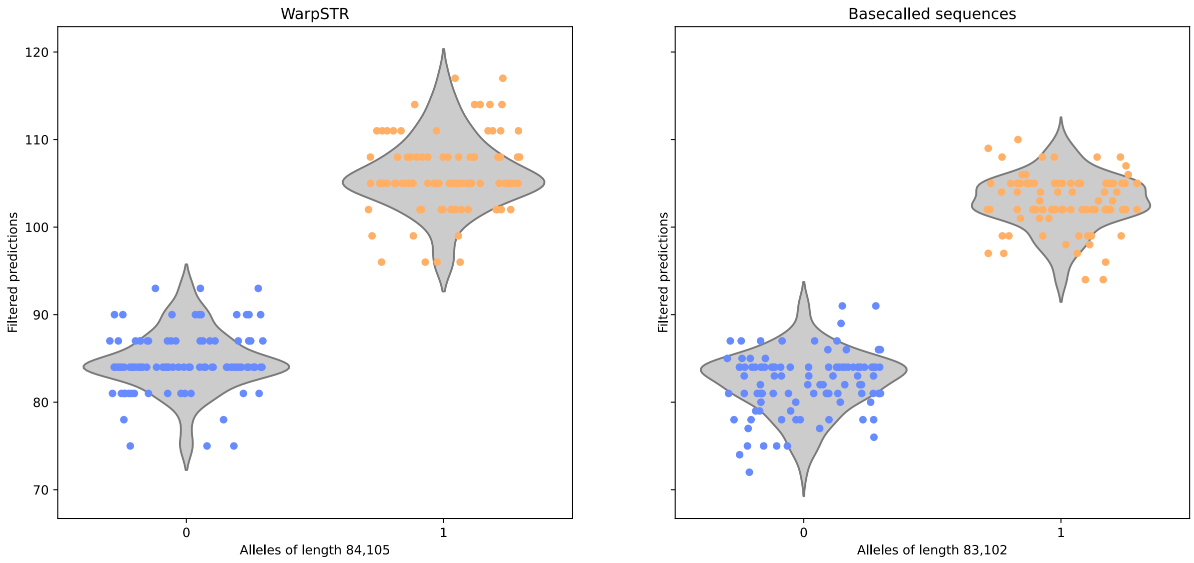


**Supplementary Figure S5:** Predictions for a whole HD locus of NA24385 subject.

### S5 Per unit predictions

In case of complex STRs, WarpSTR also predicts the number of repeat units per each repeating pattern. The number of repeats is read out from the state automaton transitions. These predictions are also genotyped using the same method based on filtering and clustering, as for simple STRs. This is useful as typically only one repeating pattern is clinically relevant, or because of the case where predictions of one repeating pattern have more dispersion thus making predictions of the whole allele length more dispersed as well.

Supplementary Figure S6 shows such results for HD locus of NA24385 subject. Our input config sequence was (AGC)AACAGCCGCCAC(CGC), WarpSTR reports counts for each repeating state (denoted in parentheses). Gold standard gives two alleles, one with 17 AGC repeats and 7 CGC repeats, and the other with 24 AGC repeats and 7 CGC repeats. Using WarpSTR, we came to the same result.

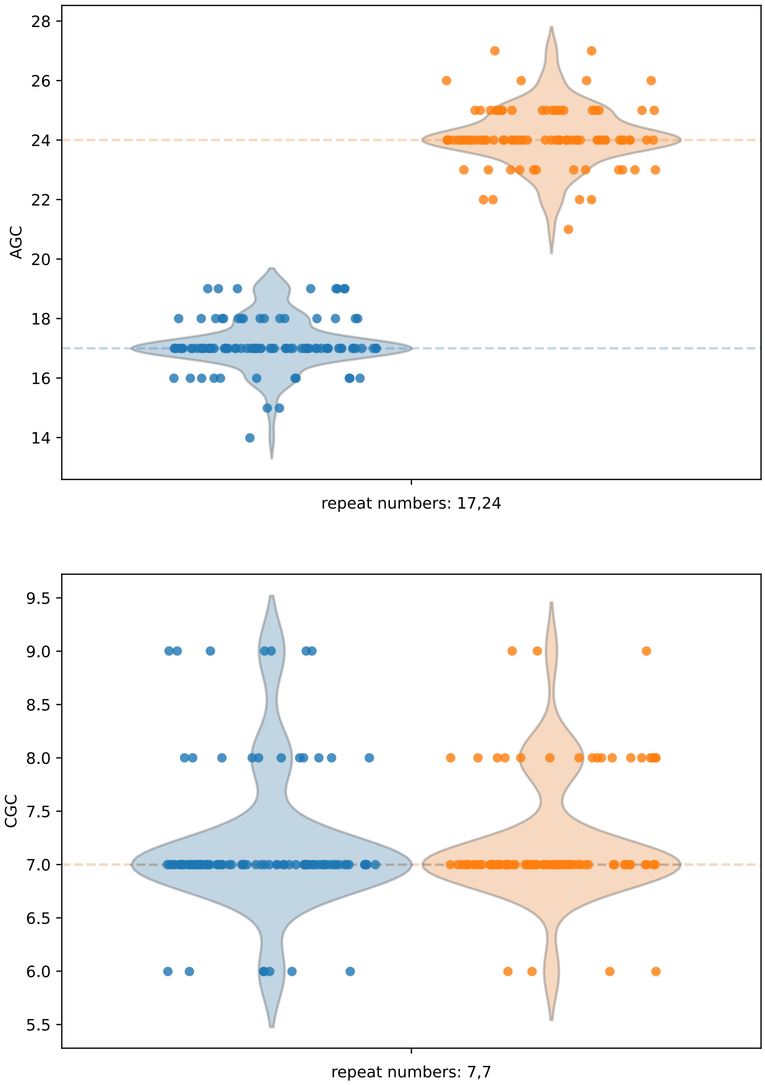


**Supplementary Figure S6:** Predictions for the HD locus of NA24385 subject as split per repeat unit.
